# 5’ Complementarity-Mediated End Joining (5’CMEJ) DNA repair

**DOI:** 10.64898/2026.07.23.740330

**Authors:** Adéla Přibylová, Cristina Ponce-Lilly, Attila Molnar, Lukáš Fischer

**Author notes:** Correspondence to: A.P., A.M., L.F. **Author contributions:** A.P.: conceptualisation, methodology, software, data curation, formal analysis, writing – original draft, visualisation, project administration. C.P.L.: investigation. A.M.: writing – review & editing, supervision. L.F.: conceptualisation, writing – review & editing, supervision, funding acquisition. A.M. and L.F. contributed equally as senior authors. All authors reviewed and approved the final manuscript.

## Abstract

How cells choose among competing DNA double-strand break repair pathways, and whether the choice differs across the tree of life, remains poorly understood, limiting prediction of CRISPR editing outcomes, particularly in plants. Through a meta-analysis of 2,098 *Sp*Cas9 target sites across plants, animals, and an alga, we show that deletion patterns reflect cell-division status rather than taxonomic origin: dividing cells favour polymerase Theta-Mediated End Joining (TMEJ), whereas non-dividing cells employ a largely overlooked pathway, 5’ Complementarity-Mediated End Joining (5’CMEJ). Unlike the known resection-dependent pathways, which use 3’ overhangs, 5’CMEJ exploits the 5’ overhangs generated by staggered *Sp*Cas9 cleavage.

## Main text

DNA double-strand breaks (DSBs) are among the most cytotoxic forms of DNA damage. They arise continuously through endogenous processes, including replication stress and oxidative metabolism, as well as from environmental insults such as ionising radiation and genotoxic chemicals. Effcient repair is therefore essential to preserve genome integrity and cellular viability across all forms of life^1^. In eukaryotes, DSBs are repaired by evolutionarily conserved pathways whose relative activity is tightly regulated throughout the cell cycle (Fig. 1a)^2,3^. In many eukaryotic cells, the dominant pathway is classical Non-Homologous End Joining (c-NHEJ), which does not require sequence homology and results in either perfect repair or short insertions and deletions (indels) when ends are processed^4^. However, when homology is available between the free DNA ends or between nearby sequences, homology-dependent pathways can be activated. The required homology length depends on the pathway, varies between organisms, and is influenced by local sequence context, amongst other factors. These pathways include polymerase Theta-Mediated End Joining (TMEJ; also known as Microhomology-Mediated End Joining, MMEJ)^5–7^, Single-Strand Annealing (SSA)^8^, and Homologous Recombination (HR)^9^. Briefly, TMEJ aligns resected 3’ ssDNA ends through short microhomologies, typically 2-6 nucleotides (nt), whereas SSA anneals longer homologous sequences within the range of tens to hundreds of nucleotides^8,10,11^. In both pathways, sequences located between the homologous regions are lost during repair, resulting in deletions at the repair junction. HR typically requires sequence homologies of hundreds to thousands of nucleotides provided *in trans* by a homologous repair template, most commonly the sister chromatid. Under these conditions, repair can proceed with high fidelity and can often result in error-free outcomes. TMEJ, SSA, and HR all rely on homologies exposed on 3’ single-stranded DNA (ssDNA) overhangs generated by MRN complex-initiated 5’ end resection^12,13^.

**Fig 1.**
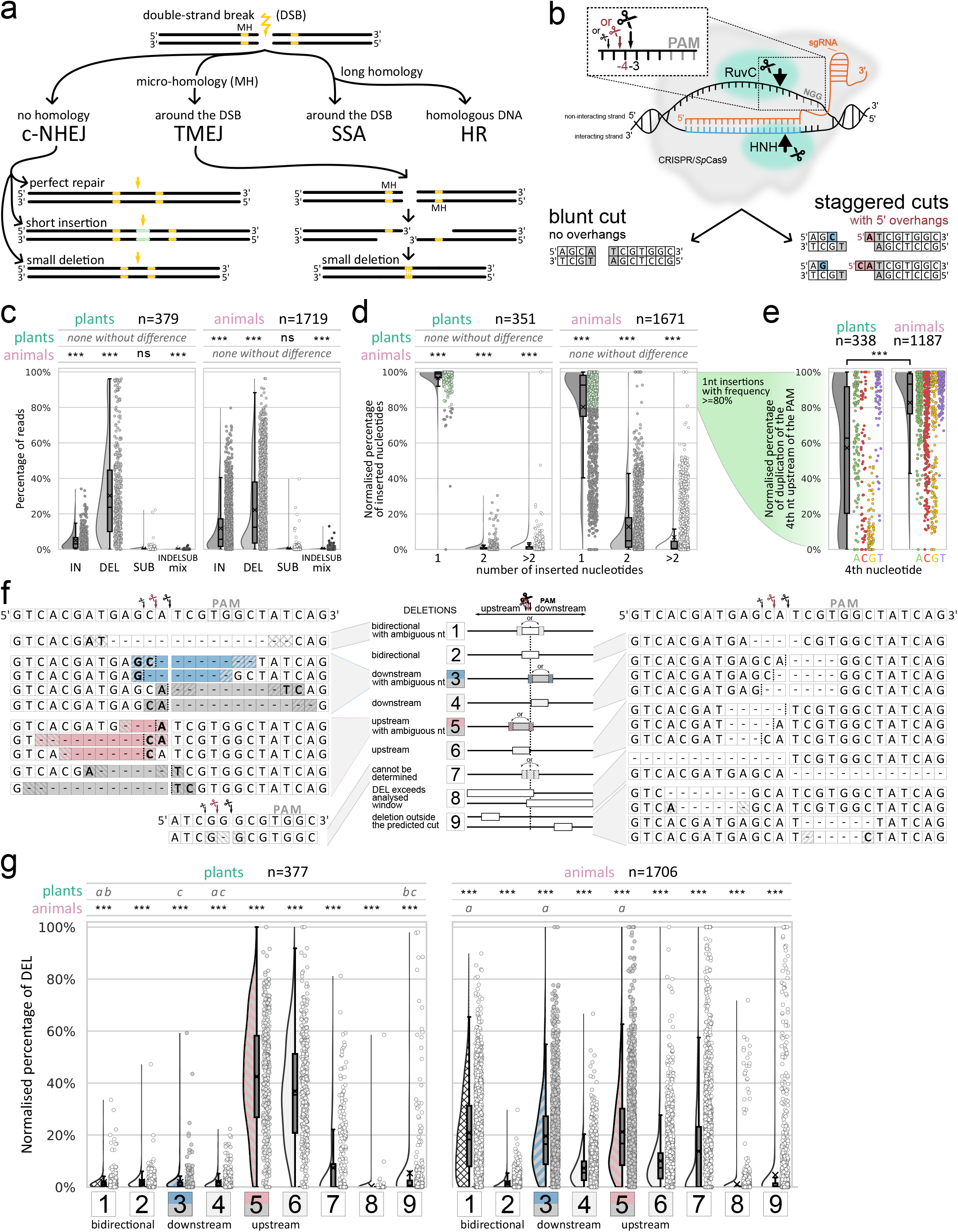
Overview of double-strand break (DSB) repair pathways at *SpCas9-induce*d breaks and comparison of mutation spectra between plant and animal datasets. **a**, Schematics of the four major DSB repair pathways: classical Non-Homologous End Joining (c-NHEJ), polymerase Theta-Mediated End Joining (TMEJ), Single-Strand Annealing (SSA), and Homologous Recombination (HR). Yellow lightning bolt indicates the DSB, yellow bars microhomology (MH) regions. **b**, Schematic of *SpCas9 cleava*ge. The sgRNA (orange) base-pairs with the interacting strand of the targeted DNA, the HNH domain cleaves it 3 nt upstream of the PAM, the RuvC domain cleaves the non-interacting strand 3–5 nt upstream of the PAM, producing either a blunt or a staggered cut with 5’ overhangs. **c**, Normalised percentage of reads assigned to insertions (IN), deletions (DEL), substitutions (SUB), and mixed indel-substitution events (INDELSUB mix) in plant (n=379) and animal (n=1,719) target sites (IN+DEL+SUB+INDELSUB mix+not mutated=100%). **d**, Normalised percentage of insertions by size (1nt+2nt+>2nt=100%) in plant (n=351) and animal (n=1,671) target sites. The green shaded region indicates sites where 1 nt insertions account for ≥80% of all insertions; such sites are further analysed in panel e. **e**, Normalised percentage of 4th-nucleotide duplication among 1 nt insertions, stratified by the identity of the 4th nucleotide upstream of the PAM (A, C, G, T), in plant (n=338) and animal (n=1,187) target sites. f, Schematic of the nine deletion categories defined by CRISPRlysis, based on position relative to the *SpCas9 cut si*te (bidirectional, downstream, upstream) and presence or absence of ambiguous nucleotide(s). Ambiguous nucleotides are highlighted on a striped background. g, Normalised percentage of deletions in each of the nine categories in plant (n=377) and animal (n=1,706) target sites (sum of categories 1 to 9 = 100%). Statistical comparisons are shown above each raincloud plot (half-violin + box plot + jittered points). Results for groups compared within a kingdom (IN plants vs DEL plants, etc.) are in the row corresponding to the tile of the violin plot; each group was compared against all other groups. Within-kingdom comparisons used the Wilcoxon signed-rank test on CLR-transformed values with Benjamini–Hochberg FDR correction; groups sharing a letter (a, b, c) are not significantly different; ‘none without difference’ indicates all groups differ. Between-kingdom comparisons (IN plants vs IN animals, etc.) used the Kruskal–Wallis test with Dunn’s post hoc test and Holm–Bonferroni correction; ns, not significant; ***p≤0.001. Each dot represents one target site; box plots show quartiles, the median (horizontal line), and the mean (×, cross). Differences in n across panels reflect restriction to sites carrying the relevant event. Full statistical details are in the Materials and Methods.

These DNA repair pathways play a central role in CRISPR/Cas-induced mutagenesis, as they process the DSBs generated by the Cas ribonucleoprotein complex, consequently determining the precision and spectrum of the editing outcomes (RNP)^14–16^. The most widely used variant, *Sp*Cas9, recognises a G-rich Protospacer Adjacent Motif (PAM; which is NGG, where N is any nucleotide), unwinds the DNA upstream of the PAM, and base-pairs with the interacting strand through its sgRNA. The interacting strand is then cleaved by the HNH domain 3 nt upstream of the PAM, while the non-interacting strand is cleaved by the RuvC domain 3-5 nt upstream of the PAM. Depending on the cut position on the non-interacting strand, the resulting DSB can be either blunt or staggered with 5’ overhangs (Fig. 1b)^12,17–19^. Because the spectrum of mutations recovered at *Sp*Cas9 cut sites reflects the underlying choice of repair pathway and end-processing events, detailed characterisation of mutation signatures can reveal previously unappreciated features of DSB repair.

Our previous work, which combined CRISPR/Cas9-induced DSBs with targeted amplicon sequencing in *Nicotiana benthamiana*, revealed striking differences in DNA repair patterns compared with those reported in animal studies^20^. These differences lie outside the scope of current CRISPR repair outcome predictors (e.g. inDelhpi, FORECasT, Lindel)^21–23^, which have been trained almost exclusively on mammalian data. We also observed deletion patterns indicating a significant involvement of a specific repair mechanism based on a homology of 5’ overhangs^20^, which we hereafter call 5’ Complementarity-Mediated End Joining (5’CMEJ), a pathway not previously described. Our original data suggest that differences in mutation patterns may be taxa- and also sample-specific.

To systematically compare DNA repair in plants and animals, and to assess 5’CMEJ occurrence, we performed a meta-analysis of Next-Generation Sequencing (NGS) data from *Sp*Cas9-targeted loci across five plant species (*Arabidopsis thaliana, Nicotiana benthamiana, Petunia hybrida, Silene latifolia*, and *Solanum lycopersicum*), one green alga (*Chlamydomonas reinhardtii*), six mammalian cell lines (HAP1, HeLa, HCT116, HEK293, Jurkat, and K562), and two animal species (*Xenopus tropicalis* and *Pleurodeles waltl*). Altogether, we analysed 379 plant and 1,719 animal target sites (individual sources listed in Supplementary Table 1^20,24–34^). To enable consistent classification of mutation patterns at *Sp*Cas9-induced DSBs across these diverse datasets, we developed CRISPRlysis, an in-house script for the systematic annotation of mutation events from amplicon sequencing data (https://github.com/pribylad/CRISPRlysis), which was validated on randomised datasets; a full description of CRISPRlysis is provided in Supplementary File 1, and its decision algorithm in Supplementary Table 2. Because the source studies differ in delivery method, cell type and sequencing depth, all comparisons below are based on sample-normalised percentages (see Materials and Methods), allowing mutation spectra in individual target sites to be compared across these heterogeneous datasets.

CRISPRlysis revealed substantial differences in mutational outcomes between plant and animal samples, confirming our previous findings in Přibylová et al., 2022^20^. Although deletions were predominant in both groups, plant samples showed relatively fewer insertions and more deletions than animal samples (Fig. 1c). Among insertions, single-nucleotide events predominated: they accounted for at least 80% of all insertions in 96% of plant and 71% of animal target sites (Fig. 1d). Longer insertions (≥2 nt) appeared to be enriched in animals and to be cell-line-dependent, occurring most frequently in the Jurkat cell line (individual sample analyses are provided in Supplementary File 2). The single-nucleotide insertions were mostly templated duplications of the 4th nucleotide upstream of the PAM, likely generated by fill-in of staggered ends and subsequent ligation, and were significantly more frequent in animals than in plants (median 93% vs 63%; Fig. 1e). As shown previously, the duplication frequency depended on the identity of the 4th nucleotide, where T and A were duplicated most often, and G the least^23,35^. G duplication, in particular, was barely detectable across plant samples relative to animal ones. Overall, the high level of the 4th-nucleotide duplication indicates frequent formation of 5’ overhangs across the majority of target sites cleaved by *Sp*Cas9.

Next, we analysed the deletion events in detail. CRISPRlysis sorted deletions into nine categories based on the position of the deletion relative to the *Sp*Cas9 cut site and on the presence of so-called ambiguous nucleotide(s) flanking the deletion boundaries. Ambiguous nucleotides are those identical on both sides of the deletion junction (typically 1-2 nt) and therefore mappable to either end. Their presence suggests that complementarity between the free DNA ends facilitated end joining (Fig. 1f); here and throughout, ‘upstream’ and ‘downstream’ denote the 5’ and 3’ sides of the cut, respectively, with the PAM on the 3’ side. Deletions spanning the *Sp*Cas9 cut site bidirectionally, or occurring downstream or upstream of it, were sorted into categories 1, 3, and 5, respectively, when ambiguous nucleotide(s) were present. When no ambiguous nucleotide(s) were detected, bidirectional, downstream, and upstream deletions were assigned to the categories 2, 4, and 6, respectively. Categories 7, 8, and 9 represent deletions whose orientation cannot be determined, that exceed the analysed window, or that occur outside the predicted cut site, respectively.

Deletion frequencies differed significantly between the plant and animal datasets across all categories (Fig. 1g), pointing to substantial differences in DNA end processing and repair. In plants, upstream deletions involving ambiguous nucleotides dominated (category 5, median 42%), followed by unambiguous upstream deletions (category 6, median 36%). In animals, by contrast, the bidirectional, downstream, and upstream categories with ambiguous nucleotides (categories 1, 3, and 5) dominated and did not differ significantly from one another (medians of 18%, 17%, and 17%, respectively). Figures 1d and 1g reveal very high dispersion among insertion and deletion data points, reflecting significant intra-sample variation of the mutation outcome across individual target sites (Supplementary File 2), suggesting that a categorisation based on kingdom alone is insuffcient.

To examine differences at the level of individual samples, defined by a unique combination of organism, experiment, and sampling time, we used Principal Component Analysis (PCA) to compare the deletion distributions across all 29 datasets analysed above, followed by *k*-means clustering of the sample coordinates in PC1-PC2 space, which resolved three clusters (A, B, and C; Fig. 2a) The first two principal components explaining 64% and 20% of the total variance, and PC1 separates samples along an upstream-versus-bidirectional/downstream axis, with upstream categories loading positively (categories 5 and 6) and bidirectional and downstream categories negatively (categories 1 and 3). PC2 mainly separates cluster B from A and C on the vertical axis because of the short 1-2 nt deletions whose orientation cannot be determined (category 7; Supplementary File 3). One cluster comprises exclusively plant samples, whereas the other two are dominated by animal samples, with some plant and algal samples interspersed. Inspecting these outliers, namely *A. thaliana* root-derived callus (in cluster B) and *C. reinhardtii* samples (in cluster A), we found that both consist of proliferating cells, which also applies to all animal samples with which these plant samples were clustered. In contrast, the remaining plant samples derived from differentiated cells formed a distinct cluster C. This grouping is not explained by the delivery method, as RNP-delivered samples are distributed across all three clusters. Likewise, *Agrobacterium*-delivered samples span both cluster B and C. Cluster identity therefore tracks cell-division status rather than the route by which *Sp*Cas9 was delivered. Consistent with this link to proliferation, human cell-line samples containing the same target sites but collected at successive time points fell into different clusters: all three lines (HCT116, HEK293, and K562) transitioned from cluster B to A between 8 and 24 hours post-RNP delivery (hpd), with K562 transitioning earliest. Transfection results in changes in gene expression and attenuated cell proliferation^36^, which may also influence the activity of individual DNA repair machineries in a time-dependent manner. These findings indicate that the repair signatures at *Sp*Cas9 breaks are shaped by cell-cycle state and cellular context, not by taxon alone, in line with the sample-specific differences noted above.

**Fig 2.**
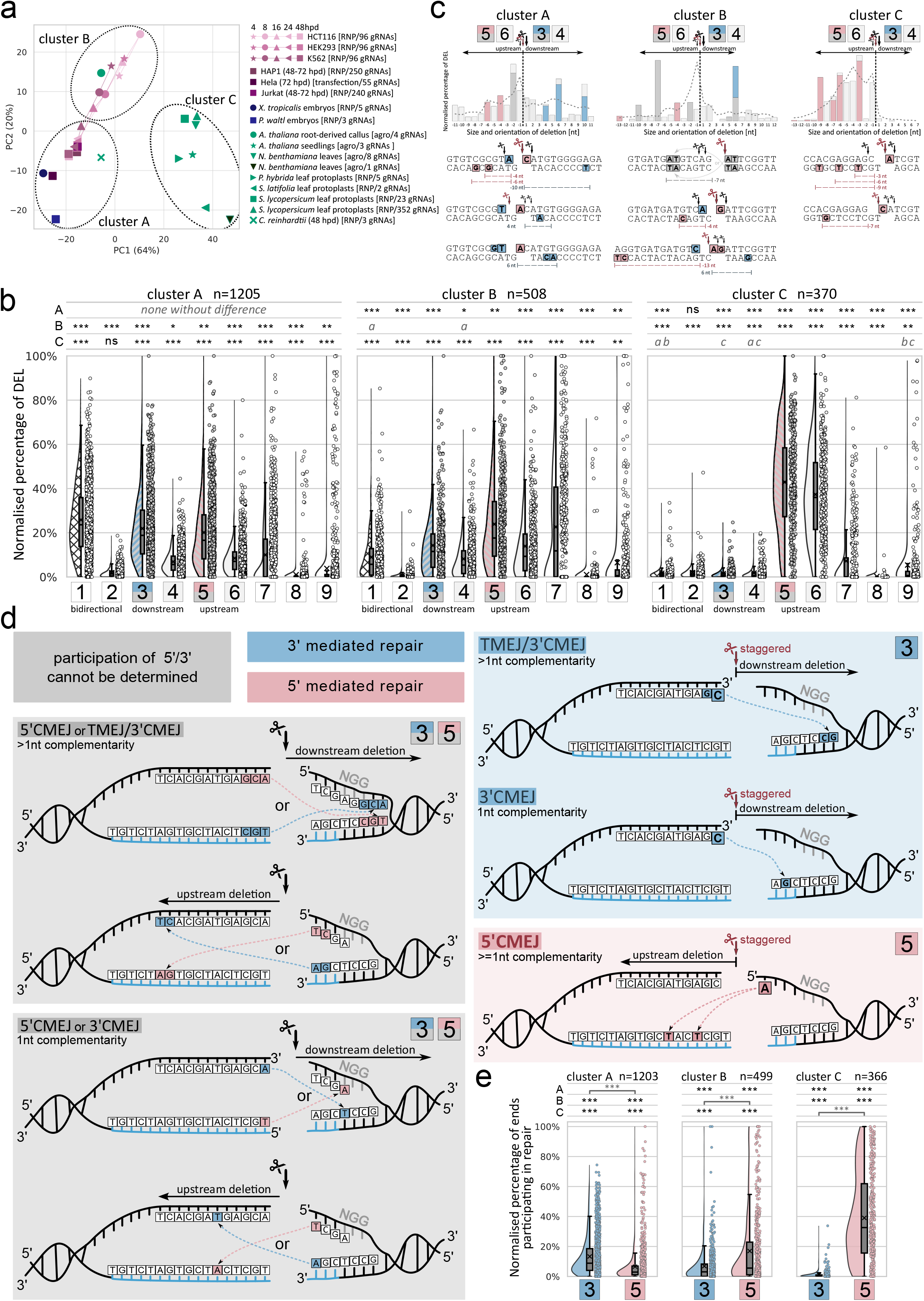
Sample-level clustering resolves a division-linked shift between 5’- and 3’-mediated end joining. **a**, Principal Component Analysis of deletion-category distributions across 29 sample datasets (key, right). PC1 and PC2 explain 64% and 20% of the total variance. Dashed ellipses mark the three clusters (A, B, C) resolved by k-means clustering of the PC1-PC2 coordinates. Dashed lines connect human cell-line samples from earlier to later timepoints (4, 8, 16, 24, 48 hours post-RNP delivery; hpd). **b**, Normalised percentage of deletions in each of the nine categories across clusters A, B, and C (n = 1,205, 508, 370 target sites; sum of categories 1-9 = 100%). **c**, Schematics of stacked bidirectional histograms of deletion size and orientation, one example per cluster. Bars left and right of the predicted cut (vertical dashed line) represent upstream and downstream deletions, respectively. A dashed-line plot represents the average distribution across all target sites in an individual cluster. Pink and blue bars represent deletions mediated by the 5’ and 3’ DNA ends, respectively; dark grey deletions are mediated by either the 5’ or 3’ end, and light grey deletions do not have complementary nucleotides. The representative alignments below illustrate examples of deletions and the nucleotides that could mediate repair. **d**, Mechanistic model. Grey panels (left), events in which 5’ and 3’ DNA end participation cannot be distinguished; blue panel (right top), TMEJ/3’CMEJ mediating downstream deletions; pink panel (right bottom), 5’CMEJ mediating upstream deletion. Highlighted nucleotides mark the complementary termini mediating each join. **e**, Normalised percentage of ends unambiguously assignable to the TMEJ/3’CMEJ (blue, subcategory of category 3) or 5’CMEJ (pink, subcategory of category 5) across clusters A, B, and C (n = 1203, 499, 366), remaining categories are in Supplementary File 2. Statistical comparisons are shown above each raincloud plot (half-violin + box plot + jittered points). Results for groups compared within a category (category 1 cluster A vs category 2 cluster A, etc.) are in the row corresponding to the tile of the violin plot; each category was compared against all other categories. Within-category comparisons used the Wilcoxon signed-rank test on CLR-transformed values with Benjamini–Hochberg FDR correction; categories sharing a letter (a, b, c) are not significantly different; ‘none without difference’ indicates all categories differ. Between-clusters comparisons (category 1 cluster A vs category 1 cluster B, etc.) used the Kruskal–Wallis test with Dunn’s post hoc test and Holm–Bonferroni correction; ns, not significant; *p≤0.05, **p≤0.01, ***p≤0.001. Each dot represents a target site; box plots show quartiles, the median (horizontal line), and the mean (×, cross). Differences in n across panels reflect restriction to sites carrying the relevant event. Full statistical details are in the Materials and Methods.

Next, we characterised cluster-specific differences in deletion spectra and examined the distribution of the deletion categories across clusters A, B, and C (n=1205, 508, and 370, respectively; Fig. 2b). In cluster A, deletions with ambiguous nucleotides (categories 1, 3, and 5) dominate across all orientations, with bidirectional deletions being most frequent, followed by downstream and upstream orientations, yielding medians of 24%, 19%, and 17%, respectively. Cluster B similarly favours deletion categories with ambiguous nucleotides over unambiguous ones. However, the balance among orientations is shifted markedly: upstream deletions are most frequent, followed by downstream and bidirectional deletions (medians 18%, 12%, and 6%, respectively), and unambiguous upstream deletions (category 6, median 9%) are more frequent than ambiguous bidirectional deletions (category 1, median 6%). Cluster C exhibits the strongest upstream bias: upstream deletions with and without ambiguous nucleotides (categories 5 and 6) reach medians of 43% and 36%, respectively, while all remaining categories have medians below 2%. Across all three clusters, deletions are dominated by categories containing ambiguous nucleotides, indicating that complementarity between the free DNA ends is a pervasive feature of repair at *Sp*Cas9 breaks. The presence or absence of ambiguous nucleotide(s) offers a provisional link to the underlying repair pathway: deletions with ambiguous nucleotides are consistent with complementarity- or microhomology-mediated end-joining, whereas deletions lacking ambiguous nucleotide(s) are more consistent with c-NHEJ.

Deletions involving ≥2 ambiguous nucleotides can be produced by TMEJ, which uses 3’ overhangs generated by 5’ end resection initiated by the MRN complex^10^. However, we also observed an enriched frequency of deletions likely resulting from a repair that used only a single complementary nucleotide, which the canonical TMEJ mechanism cannot account for. Both a 3’ and a 5’ terminal nucleotide, exposed by the *Sp*Cas9 cut, can pair with a complementary nucleotide on the opposing strand to facilitate DSB repair (Fig. 2c, 2d). Pairing of the exposed 3’ terminal nucleotide of the non-interacting strand (highlighted in blue) removes sequence downstream of the cut, producing a downstream deletion with ambiguous nucleotides (category 3). In case of single-nucleotide complementarity, we refer to this route as 3’ complementarity-mediated end joining (3’CMEJ), whereas when the complementarity extends beyond 1 nt, the same event is equally consistent with TMEJ (Fig. 2d). Pairing of the exposed 5’ terminal nucleotide of the non-interacting strand (highlighted in red) instead removes sequence on the upstream side, producing an upstream deletion with ambiguous nucleotides (category 5; Fig. 2d). We term this 5’ complementarity-mediated end joining (5’CMEJ), a mechanism identified in *N. benthamiana* and described in Přibylová et al., 2022^20^. In most cases, 3’CMEJ and 5’CMEJ cannot be unequivocally distinguished in NGS data, as the deletion can often be explained by the participation of both 3’ and 5’ terminal nucleotides. These cases involve all deletions with an ambiguous nucleotide at the 3rd and 4th nucleotide upstream of the PAM for upstream and downstream deletions, respectively (Fig. 2d, left grey panels). However, *Sp*Cas9 frequently forms 5’ staggered ends, inferred from 4th-nucleotide duplication (Fig. 1e). Upstream deletions with an ambiguous nucleotide at the 4th position upstream of PAM on the non-interacting strand clearly fall into the 5’CMEJ category, as there is no 3’ nucleotide on the bottom interacting strand, which could mediate such repair (Fig. 2d red panel). Partially, the same applies to downstream deletions with ambiguous nucleotides after the staggered cut, where TMEJ/3’CMEJ mediates repair (Fig. 2d, blue panel). Importantly, this TMEJ/3’CMEJ type of repair can also happen after the formation of a blunt cut, if the 3’ terminal nucleotide/s is removed by the post-cleavage activity of the RuvC domain. The detailed description of the classification criteria for individual categories is presented in Supplementary File 1 and Supplementary Table 2. Collectively, 5’CMEJ was identifiable in all three PCA clusters (Fig. 2c), indicating that it operates across the plant, algal, and animal lineages examined here, not only in the *N. benthamiana* dataset first identified in our work shown in Přibylová et al., 2022^20^. Thus, alongside canonical TMEJ, complementarity-mediated end joining, and 5’CMEJ in particular, are recurrent contributors to deletion repair at *Sp*Cas9 breaks in both plants and animals.

To quantify the relative contribution of 5’CMEJ and TMEJ/3’CMEJ at the sample level, we compared their normalised representation across clusters A, B, and C (Fig. 2e). As a control, we also generated a dataset of 3×10,000,000 random sequences with randomised deletion events around the cut site (Supplementary File 1 and Supplementary Table 3) to assess whether the observed deletion pattern in the clusters conforms to a standard distribution and to determine the bias of 5’CMEJ and 3’CMEJ ratios caused by the presence of 5’ overhangs generated by *Sp*Cas9. In cluster A, TMEJ/3’CMEJ exceeded 5’CMEJ (medians of 8.6% and 2.0%, respectively). Relative to the corresponding ambiguous deletion pools in this cluster (categories 3 and 5 with medians of 19% and 17%, respectively; Fig. 2b), the single-nucleotide 5’ route accounted for only a small fraction of upstream ambiguous deletions, whereas the 3’ route accounted for a larger proportion of downstream ones. In cluster B, the balance shifted, with 5’CMEJ exceeding TMEJ/3’CMEJ (medians of 5.6% and 3.1%, respectively), consistent with the intermediate character of this cluster between clusters A and C. In cluster C, 5’CMEJ strongly dominated over TMEJ/3’CMEJ (medians of 32.6% and 0.3%, respectively), consistent with the upstream deletion bias in this cluster. Notably, analysis of the randomised dataset shows that there is only a 2.2% higher likelihood of observing 5’CMEJ compared to 3’CMEJ due to the nature of *Sp*Cas9 forming 5’ staggered overhangs. This difference is negligible compared to the ratios in clusters A, B, and C, where the 5’/3’CMEJ ratios are 50.8%, 28.7%, and 98.2%, respectively (Supplementary Table 3). Together, these results show that the balance between 5’CMEJ and TMEJ/3’CMEJ shifts systematically with cluster identity, and hence with cell-cycle and proliferation state: non-proliferating cells (cluster C) favour 5’CMEJ, whereas actively dividing cells (cluster A) favour TMEJ/3’CMEJ, the latter consistent with the resection-dependent, cell-cycle-linked activity of TMEJ. The presence of 5’CMEJ in all three clusters further indicates that it is not restricted to a single organism or lineage.

Taken together, our observations suggest that 5’CMEJ is not restricted to DNA repair in plants but is a previously overlooked component of the DSB repair landscape that becomes visible when resection is limited and 5’ overhangs persist, conditions expected under G0/G1 arrest and, potentially, prolonged *Sp*Cas9 retention at the cut site. When 5’-end resection is restricted, the canonical TMEJ route is disfavoured, and the exposed 5’ overhang, rather than being processed away, can itself be involved in template joining. This reframes how mutation spectra at *Sp*Cas9 breaks are interpreted: the 5’ overhang, previously regarded as a transient intermediate en route to fill-in or resection, is itself a functional repair substrate whose use is gated by cell-cycle stage, end-processing kinetics, and the post-cleavage behaviour of *Sp*Cas9. It also helps explain why deletion spectra from differentiated plant tissues differ from those of proliferating mammalian cell lines and have restricted prediction by existing outcome-prediction tools, which were trained predominantly on proliferating human cell lines in which resection is active, 5’ overhangs are processed, and 5’CMEJ plays only a minor role. Incorporating 5’CMEJ into mechanistic models of *Sp*Cas9 repair is therefore likely necessary to extend reliable outcome prediction to plants and to non-dividing cell types, with direct implications for the safety and precision of CRISPR-based applications in both gene therapy and crop breeding. Further work is needed to characterise the prevalence of 5’CMEJ across diverse eukaryotes and to identify the molecular factors that stabilise the 5’ overhang and mediate its ligation.

## Materials and Methods

### Next-generation Sequencing Data Analysis

NGS datasets were obtained from published works on plants and animals, which are listed in Supplementary Table 1. Together, 379 plant and 1,719 animal target sites were analysed. NGS data were analysed using a decision algorithm embedded in an in-house Python script, CRISPRlysis, available on GitHub (https://github.com/pribylad/CRISPRlysis); a brief description of CRISPRlysis is provided in Supplementary File 1 and the decision algorithm in Supplementary Table 2. Unless stated otherwise, each analysis presented in this work is restricted to the target sites carrying the relevant event, so sample sizes vary between panels.

### Statistical information

Samples that were compared within a single normalised dataset group were treated as compositional data that sum to 100% (samples within the plant, animal, cluster A, cluster B, and cluster C datasets; Fig. 1c,d,g, Fig. 2b,e). Prior to statistical testing, values were transformed using the centred log-ratio (CLR) transformation with a pseudocount of half the minimum non-zero value for zero-replacement. Then, pairwise differences between all mutation categories were assessed using the Wilcoxon signed-rank test (p < 0.05; Python scipy.stats.wilcoxon version: 1.16.3) on CLR-transformed values and multiple pairwise comparisons were corrected for multiple testing using the Benjamini-Hochberg FDR correction (Python statsmodels.stats.multitest.multipletests version: 0.14.6).

When samples between different groups were compared (plants vs animals, cluster A vs B vs C; Fig. 1c,d,e,g, Fig. 2b,e), normality of the data was tested by the Shapiro-Wilk test (p < 0.05; Python scipy.stats.shapiro version: 1.16.3). Data were not normally distributed and were tested by a non-parametric, the Kruskal-Wallis test (p < 0.05; Python scipy.stats.kruskal version: 1.16.3), followed by Dunn’s post hoc test with Holm-Bonferroni p-value adjustment for multiple comparisons (p < 0.05; Python scikit_posthocs.posthoc_dunn version: 0.13.0).

Principal Component Analysis (PCA; Python scikit-learn version 1.6.1) was used for multivariate analysis of deletion spectra across the 29 samples. Deletion categories 1-8 were used; category 9 (deletions outside the predicted cut site) was excluded as such events do not report on repair at the *Sp*Cas9-induced break, and remaining categories were renormalised to sum to 100% per sample. Because the input values were sample-normalised percentages on a common scale, no further standardisation was applied. Samples were partitioned by k-means clustering (Python sklearn.cluster.KMeans, k = 3, n_init = 50, random_state = 42) on the PC1-PC2 coordinates. The number of clusters was chosen from the within-cluster sum of squares, which fell sharply from k = 2 to k = 3 (6195 to 2342) with only modest further reduction thereafter, supported by silhouette analysis over k = 2-6 (maximum at k = 3). Cluster assignment was identical across 50 random initialisations (mean adjusted Rand index = 1.00; Supplementary File 3).

## Data Availability

NGS data from the analysed studies are available in free-access repositories; a list of these repositories can be found in Supplementary Table 1. *Silene latifolia* and *Chlamydomonas reinhardtii* data are available upon request.

## Code Availability

The script used to analyse the data, CRISPRlysis, is available on GitHub (https://github.com/pribylad/CRISPRlysis).

## Acknowledgements

This work was supported by the project TowArds Next GENeration Crops [reg. no. CZ.02.01.01/00/22_008/0004581] of the ERDF Programme Johannes Amos Comenius.

## Ethics declarations

Not applicable.

## Supplementary Information

Provided as a separate file.

## Licence

This work is licensed under a Creative Commons Attribution license (CC-BY 4.0 International).

