## Supplementary material for "5’ Complementarity-Mediated End Joining (5’CMEJ) DNA repair": Supplementary_Fig_1.pdf

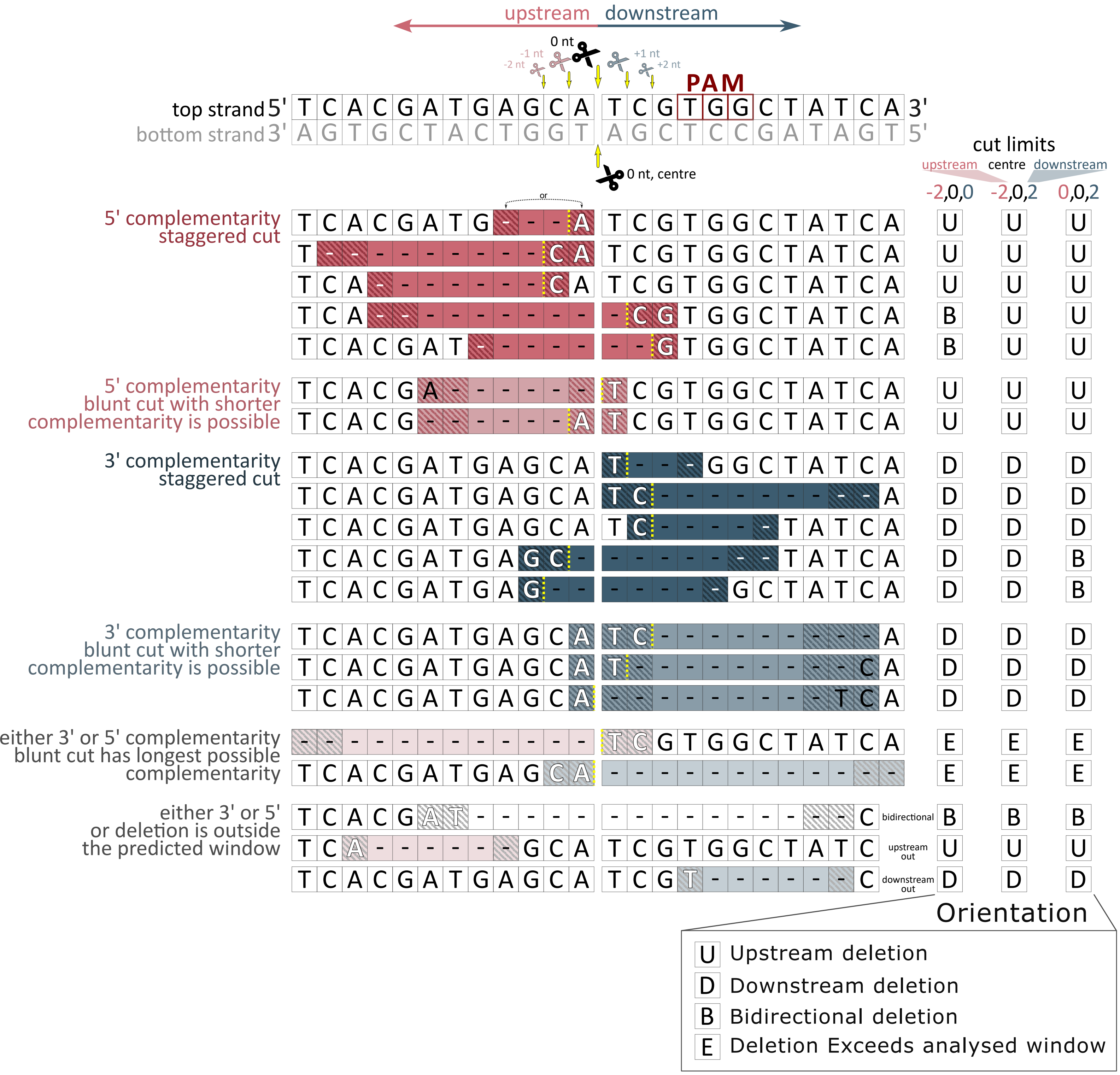


**Supplementary Fig.1|Assignment of deletion orientation and 3′/5′ DNA end participation relative to the *Sp*Cas9 cut site.**
Schematic visualisation of the target site (top and bottom strands, PAM in red) showing the canonical blunt cut at position 0 and the staggered cuts from -2 to +2 nt considered here. Upstream (red) and downstream (blue) orientations are defined relative to the cut site position. Each row below shows an example of a deletion as reported by Next-Generation Sequencing with visualisation of the top strand only. - Dashes denote deleted bases, the yellow dotted line marks the cut position, and N hatched boxes denote the ambiguous nucleotides (complementary bases) that could have mediated the repair event and can be mapped to either side of the deletion. Rows are grouped by category of deletion, and only deletions with ambiguous nucleotides are shown. The three right-hand columns give the orientation of the deletion assigned to each deletion example under three alternative settings of the permitted cut window in the CRISPRlysis software (cut limits, given as upstream, centre and downstream bounds in nt): −2,0,0; −2,0,2; 0,0,2. Orientation categories are upstream (U), downstream (D), bidirectional (B) and deletions exceeding the analysed window (E). A detailed description of deletion categories is provided in Supplementary Table 2 and the CRISPRlysis code (https://github.com/pribylad/CRISPRlysis).
