## Supplementary material for "5’ Complementarity-Mediated End Joining (5’CMEJ) DNA repair": Supplementary_File_1.pdf

### CRISPRlysis - brief description and output examples

NGS data were first analysed with CRISPResso2 (version 2.2.11)[1] using default settings, except for offset_around_cut_to_plot, which was set to the widest window for each sample set to capture the longest deletion events. The resulting Alleles_frequency_table_*.txt files were reanalysed using an in-house developed Python script, **CRISPRlysis**, available on GitHub (https://github.com/pribylad/CRISPRlysis).

In brief, the script makes a complex analysis of mutation events. Analyse frequencies and ratios of insertion, deletion, and substitution events around the cut site. For insertions, the number of inserted nucleotides and, in the case of single-nucleotide insertions, whether they are templated by the 4th nucleotide upstream of the PAM. For deletions, it analyses the ratio of deletions that are upstream or downstream of the predicted cut site, or that span the cut site. Then, the ratio of possible blunt and staggered cut formation and how they contribute to deletions upstream and downstream of the cut, with the effect of the 4th nucleotide upstream of the PAM sequence. Last but not least, the analysis of which 5'/3' DNA end is likely to participate in the repair. An example of the output is provided below and in the example analysis on GitHub. The CRISPRlysis code can be used to analyse any CRISPResso2 (version 2.2.11) Alleles_frequency_table_*.txt output file or file with the same structure. Moreover, it can analyse any type of cut (*Sp*Cas9, *Lb*Cas12a, ...); only the nuclease and cut positions must be modified in the parameters.txt file. The code's default graphical outputs were optimised for *Sp*Cas9 but can be modified for other endonucleases by changing the script parameters; see the GitHub descriptions. In this study, we set the default parameters to use reads that appear at least 5 times in the individual library. If biological replicates were available, we averaged them. We used target sites at which mutagenesis efficacy exceeded 5%.

The code was validated on a randomised dataset of 3x10,000,000 random 100 bp long sequences with random position of deletions around the cut site (middle of the sequence, position 0), where deletion starts or ends in the range -4 to +4 nucleotides from the position 0 (----|----), position 0 is in the case of *Sp*Cas9 three nucleotides upstream of the PAM sequence for the blunt type of the cut. The random datasets were tested with settings for *Sp*Cas9 (-2,0,0 -> allowing up to 2nt 5' overhang), its mirror setup (0,0,2 -> allowing up to 2nt 3' overhang), and a symmetrical combination (-2,0,2 -> allowing both types of overhang). Graphical representations of all possible cut types and their resulting deletion categories are presented in Supplementary Fig. 1 and Supplementary Table 2. The dataset was tested with the presence of the canonical PAM sequence (NGG) and without it (NNN), and PAM presence has almost no effect on the ratios of 5'/3' DNA ends possibly participating in the complementarity-mediated repair. More importantly, there is only a slight bias toward the *Sp*Cas9 type of cut, as it, by its nature, generates a -2,0,0 type cut. In the randomised dataset, there is a 2.2% higher chance of having 5' mediated repair than 3' (Supplementary Table 3). All data presented in this paper are analysed with the settings -2,0,0 unless otherwise stated; cutting in this range is the most frequent[2].

### Example of outputs

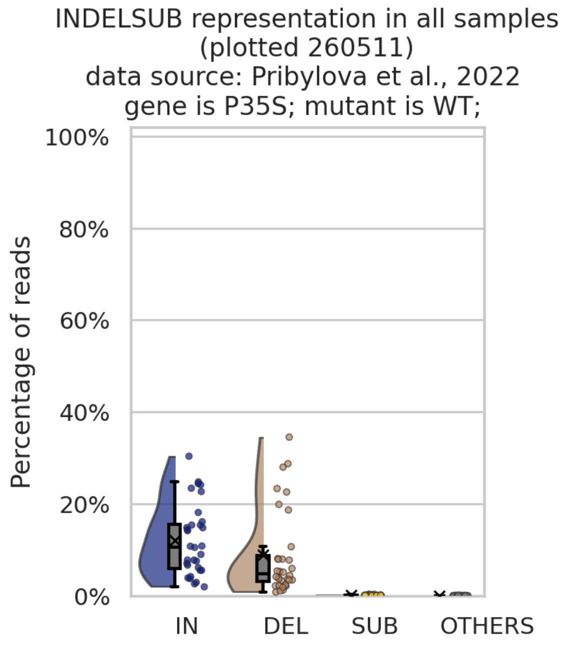


INDELSUBOTHERS category clasified only for events around the cut.
OTHERS are combinations of INs, DELs and SUBs (INDELs etc.).
IN+DEL+SUB+OTHERS+no_mutation = 100%.
Analysed 8 gRNAs in 4 groups.

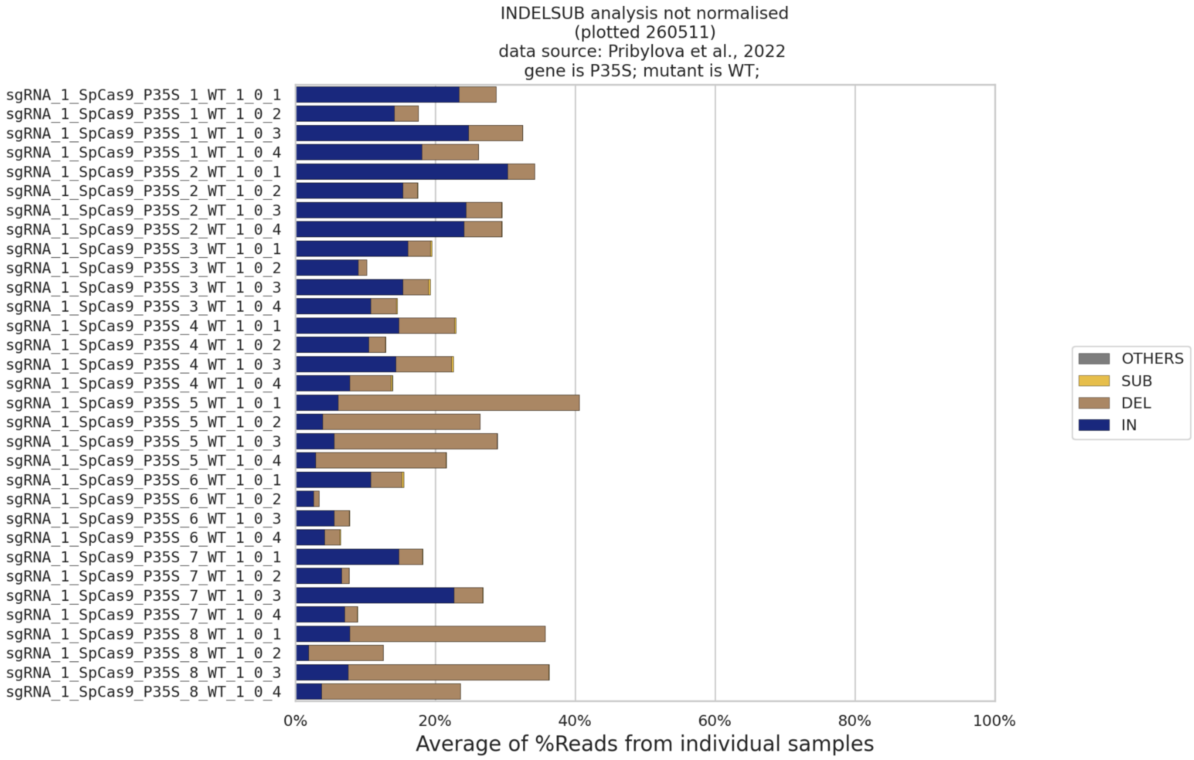


INDELSUBOTHERS category clasified only for events around the cut.
OTHERS are combinations of INs, DELs and SUBs (INDELs etc.).
IN+DEL+SUB+OTHERS+no_mutation = 100%.

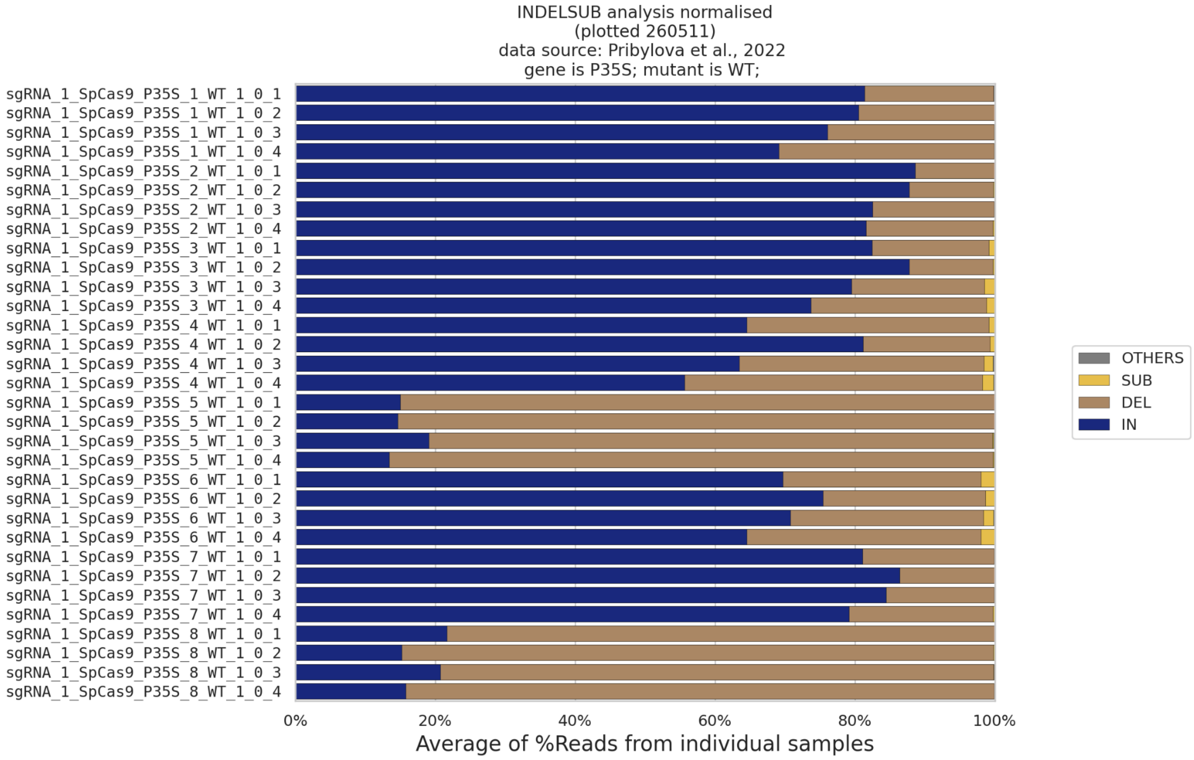


INDELSUBOTHERS category clasified only for events around the cut.
OTHERS are combinations of INs, DELs and SUBs (INDELs etc.).
IN+DEL+SUB+OTHERS = 100%.

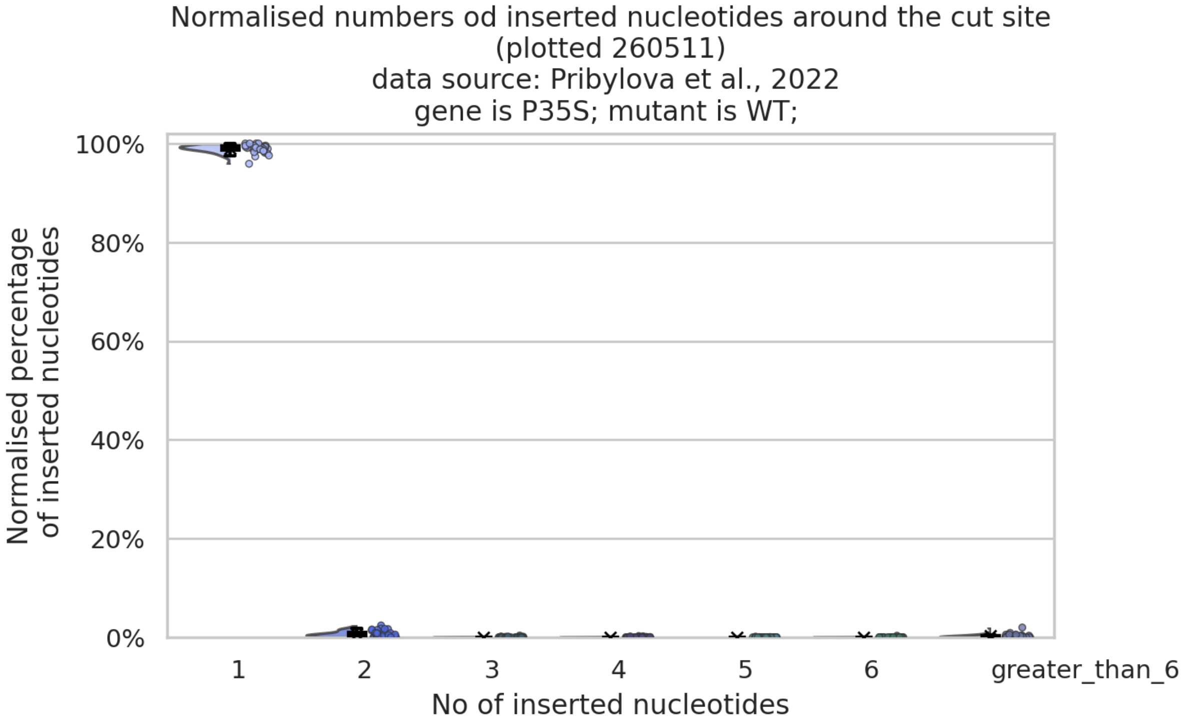


Only IN events around cut.
Sum of all IN in one sample = 100%
Analysed 8 gRNAs in 4 groups.

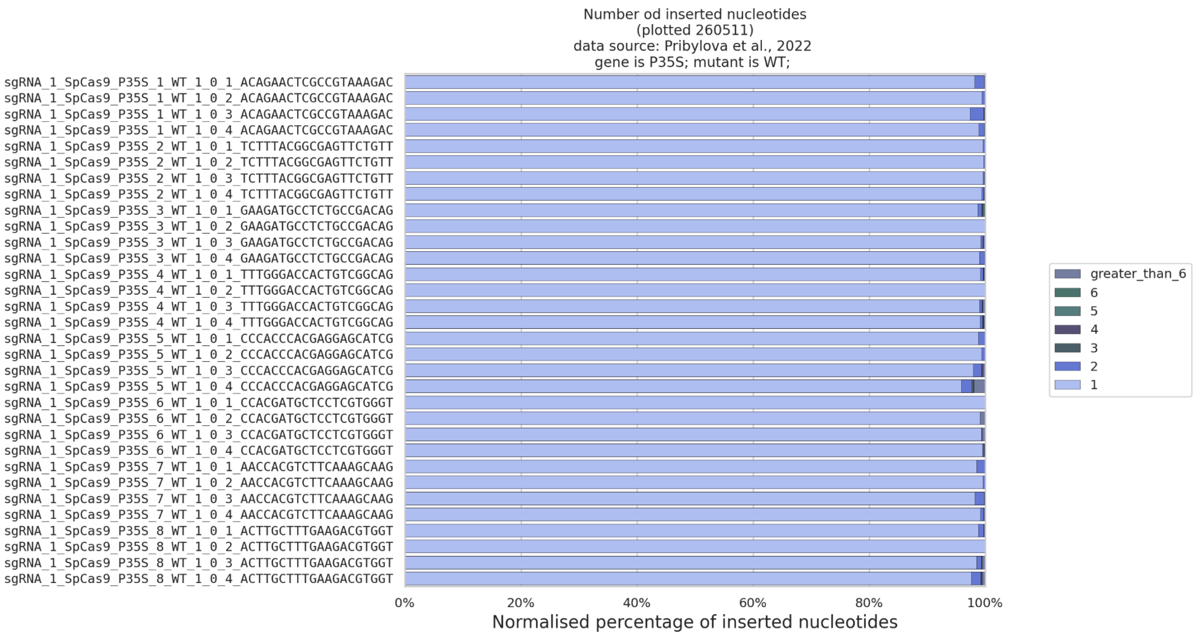


Only IN events around cut.
Sum of all IN in one sample = 100%
Analysed 8 gRNAs in 4 groups.

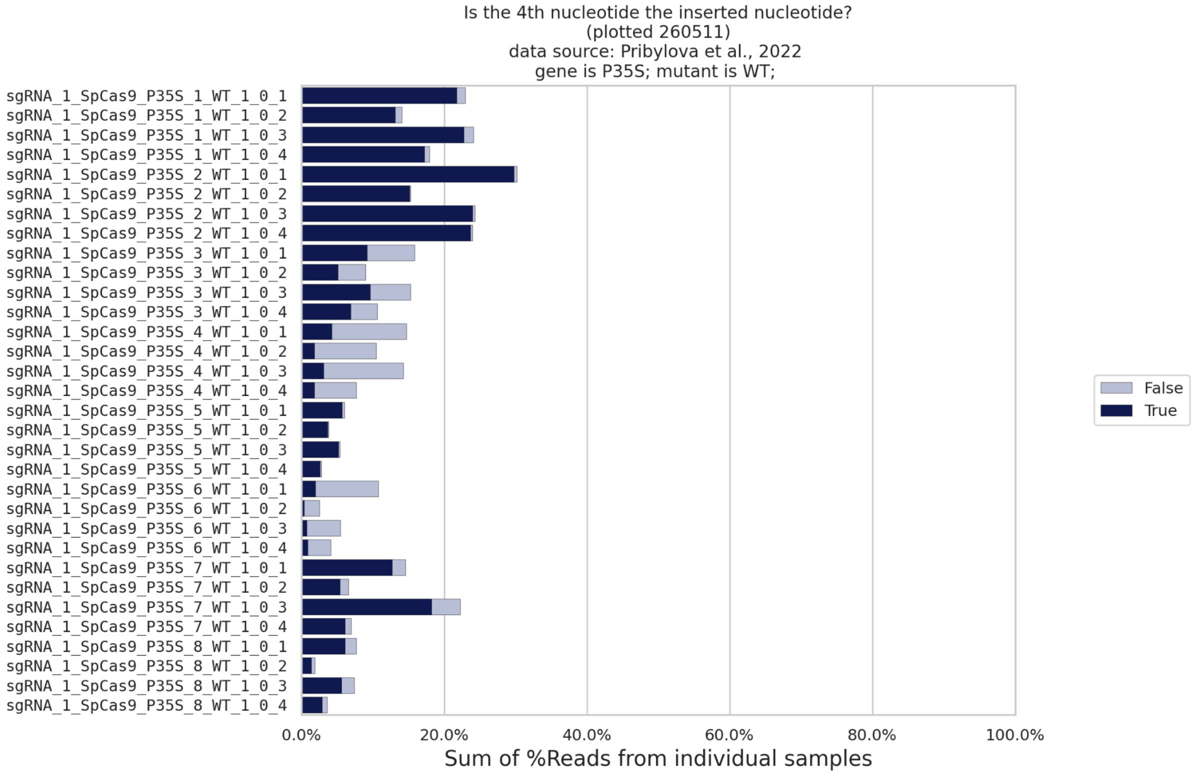


Only 1nt IN around the cut are plotted.
Sequences without DEL/SUB around the cut.

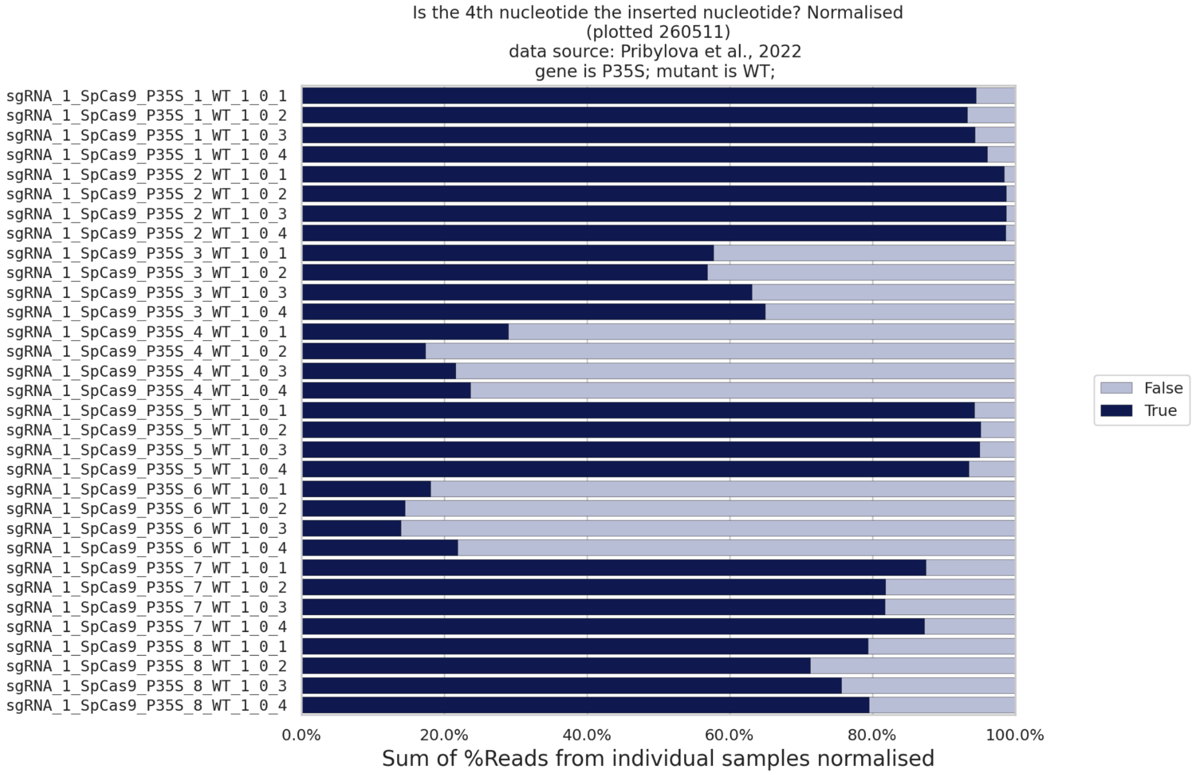


Only 1nt IN around the cut are plotted.
Sequences without DEL/SUB around the cut.

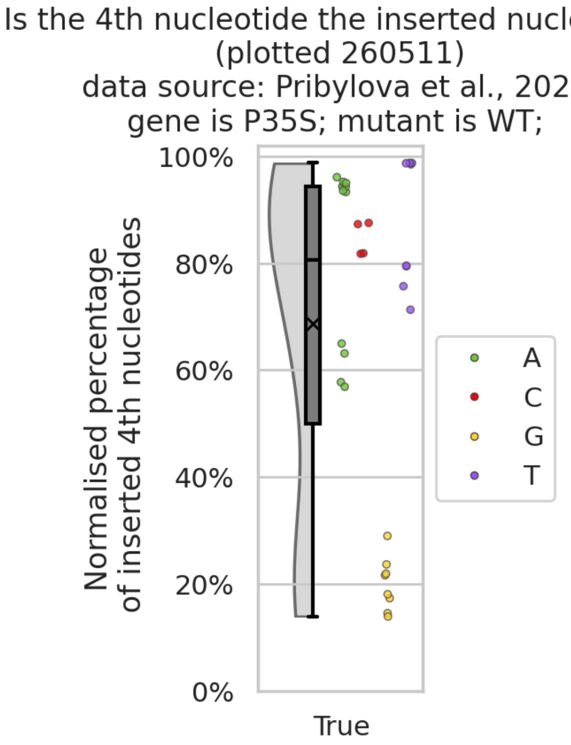


Only 1nt IN around the cut are plotted.
Sequences without DEL/SUB around the cut.
In individual sample True + False = 100%.
Analysed 8 gRNAs in 4 groups.

Is the 4th nucleotide the inserted nucleotide?
(plotted 260511)
data source: Pribylova et al., 2022
gene is P35S; mutant is WT;

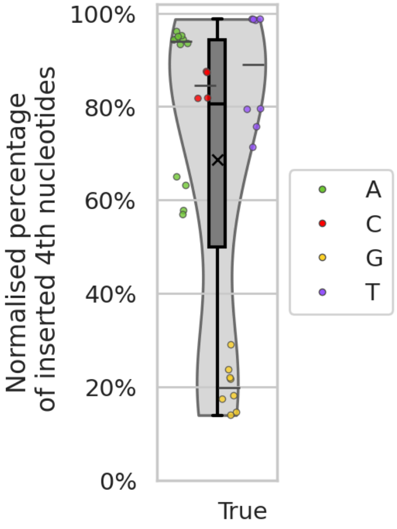


Only 1nt IN around the cut are plotted.
Sequences without DEL/SUB around the cut.
In individual sample True + False = 100%.
Horizontal lines show medians for individual 4th nucleotide.
Analysed 8 gRNAs in 4 groups.

Not-normalised incidence of DEL according to the orientation
(plotted 260511)
data source: Pribylova et al., 2022
gene is P35S; mutant is WT;

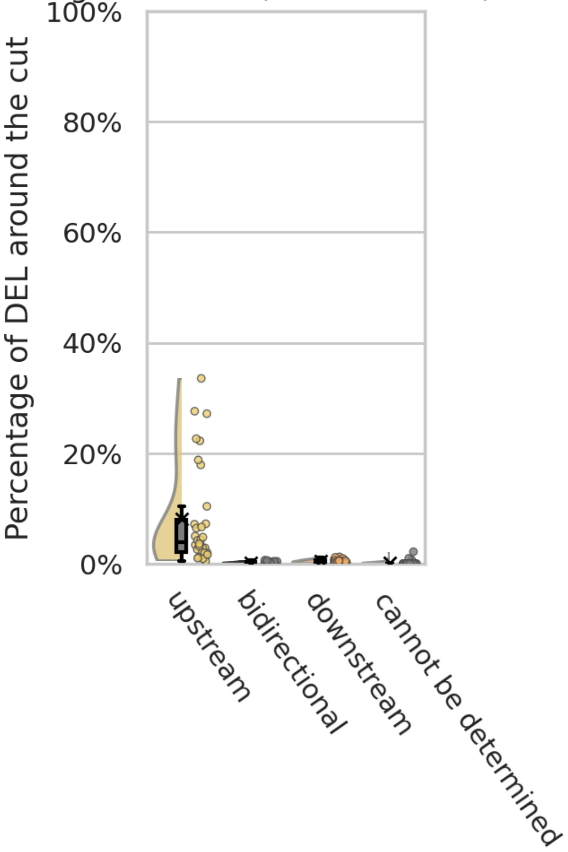


Only DELs around the cut are plotted.
Not-normalised
Analysed 8 gRNAs in 4 groups.

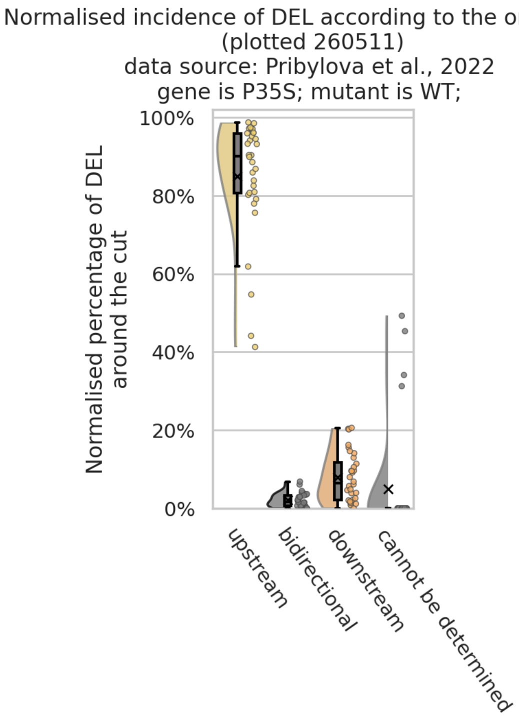


Only DELs around the cut are plotted.
Normalised to: SUM of all DEL orientation in individual sample = 100%
Analysed 8 gRNAs in 4 groups.

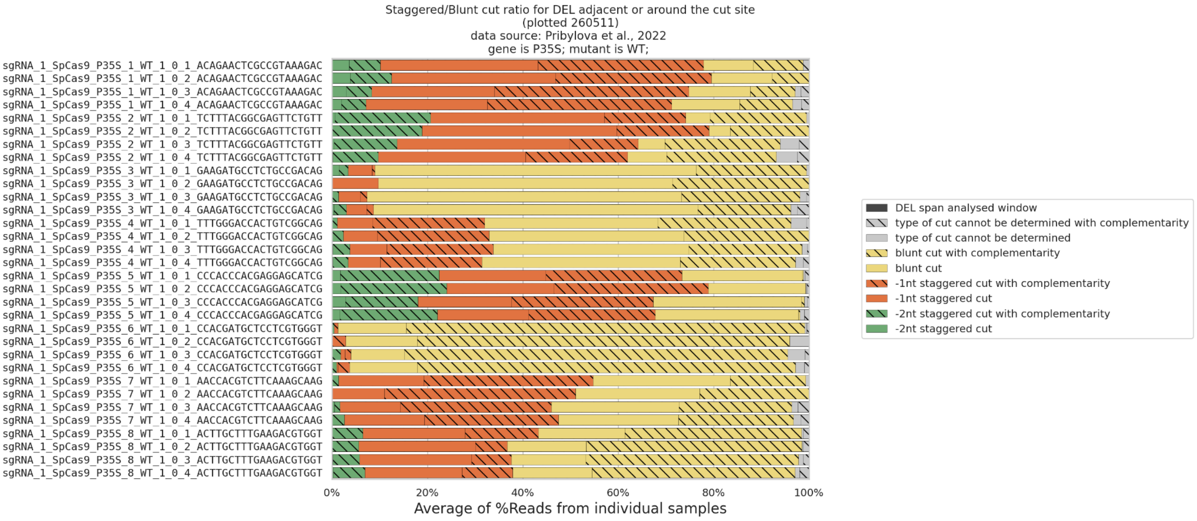


Only DELs around the cut are plotted.
Normalised to: SUM of all DEL categories around the cut in individual sample = 100%.

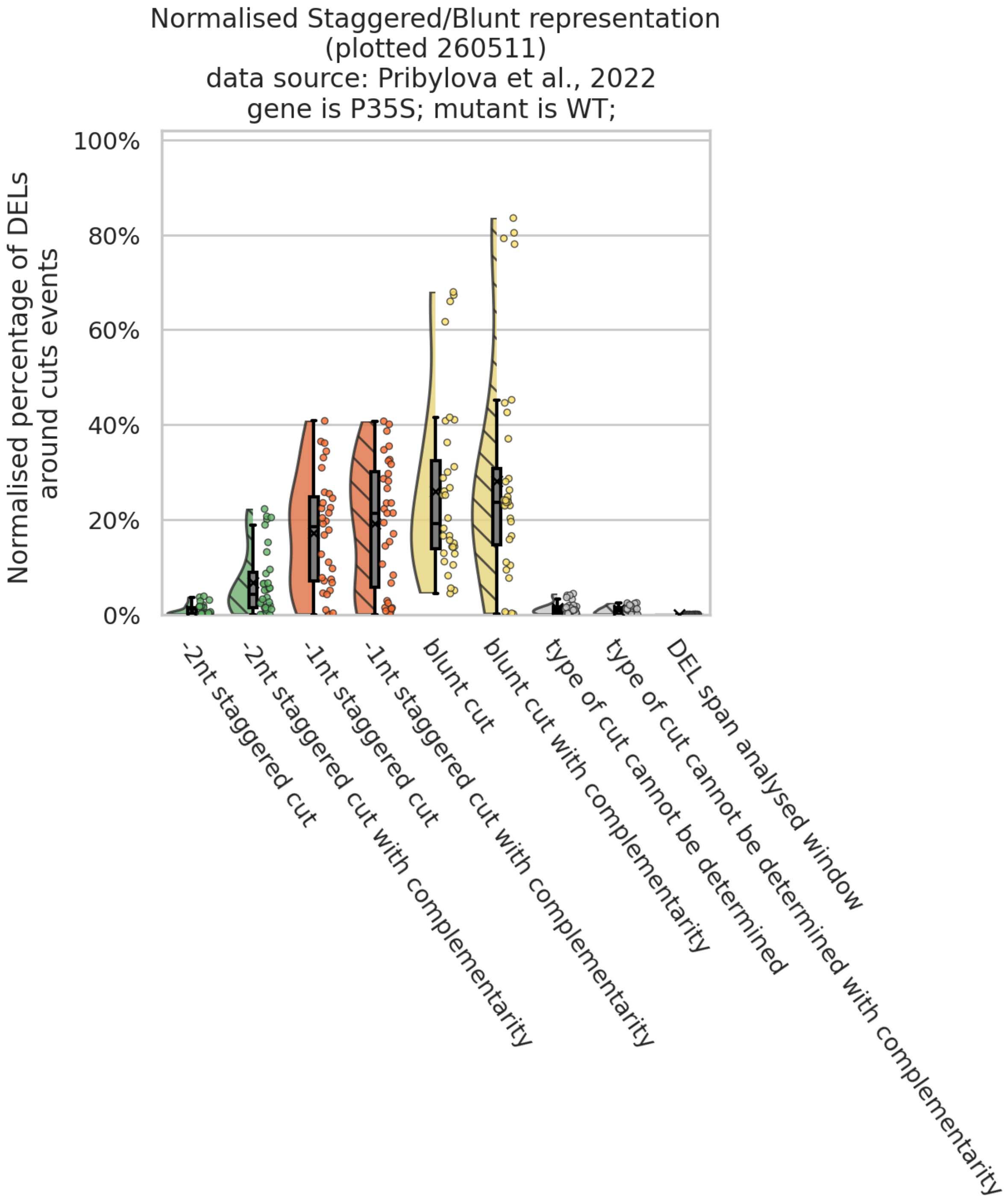


Only DELs around the cut are plotted.
Normalised to: SUM of all DEL categories around the cut
in individual sample = 100%.
Analysed 8 gRNAs in 4 groups.

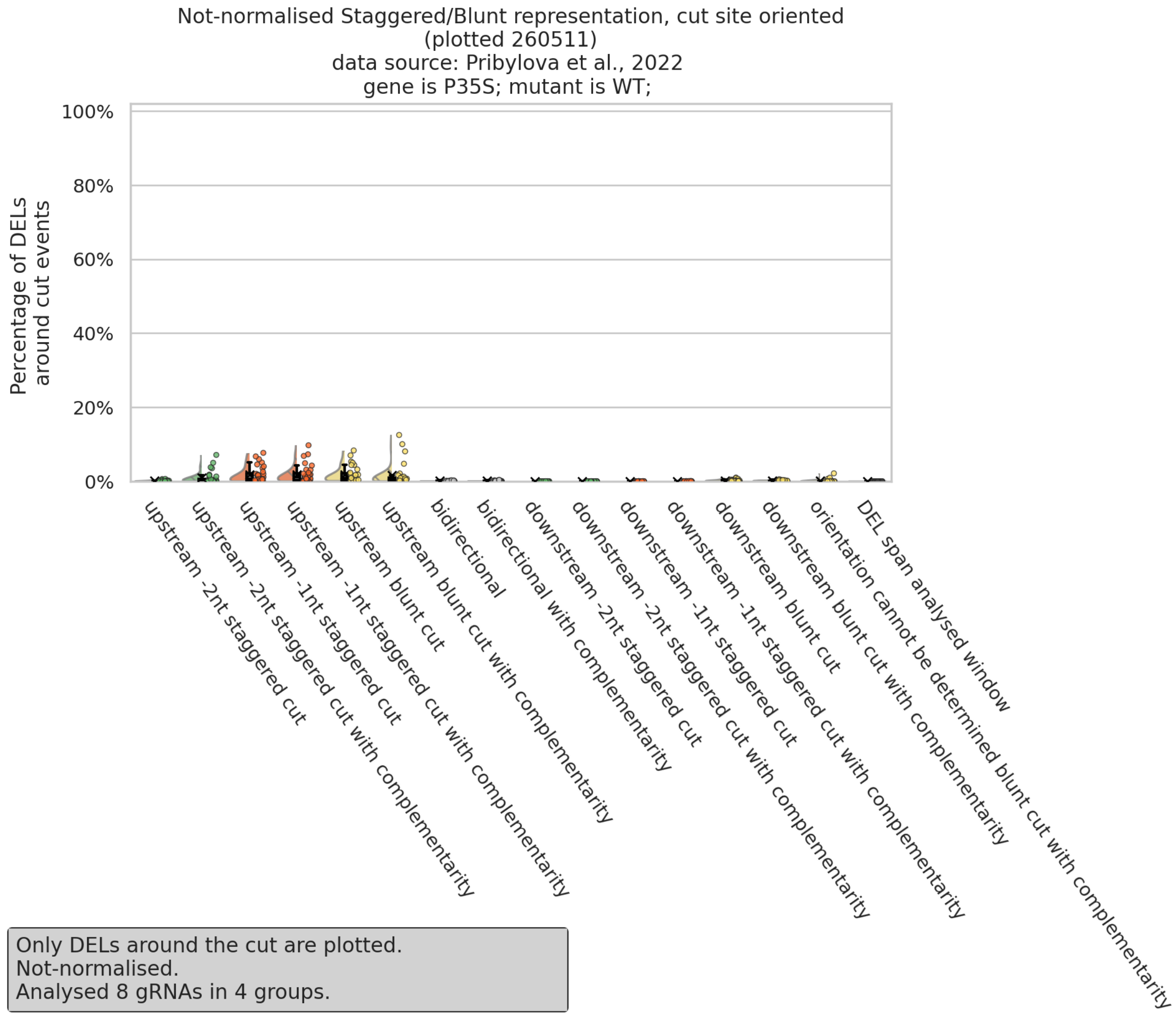


Only DELs around the cut are plotted.
Not-normalised.
Analysed 8 gRNAs in 4 groups.

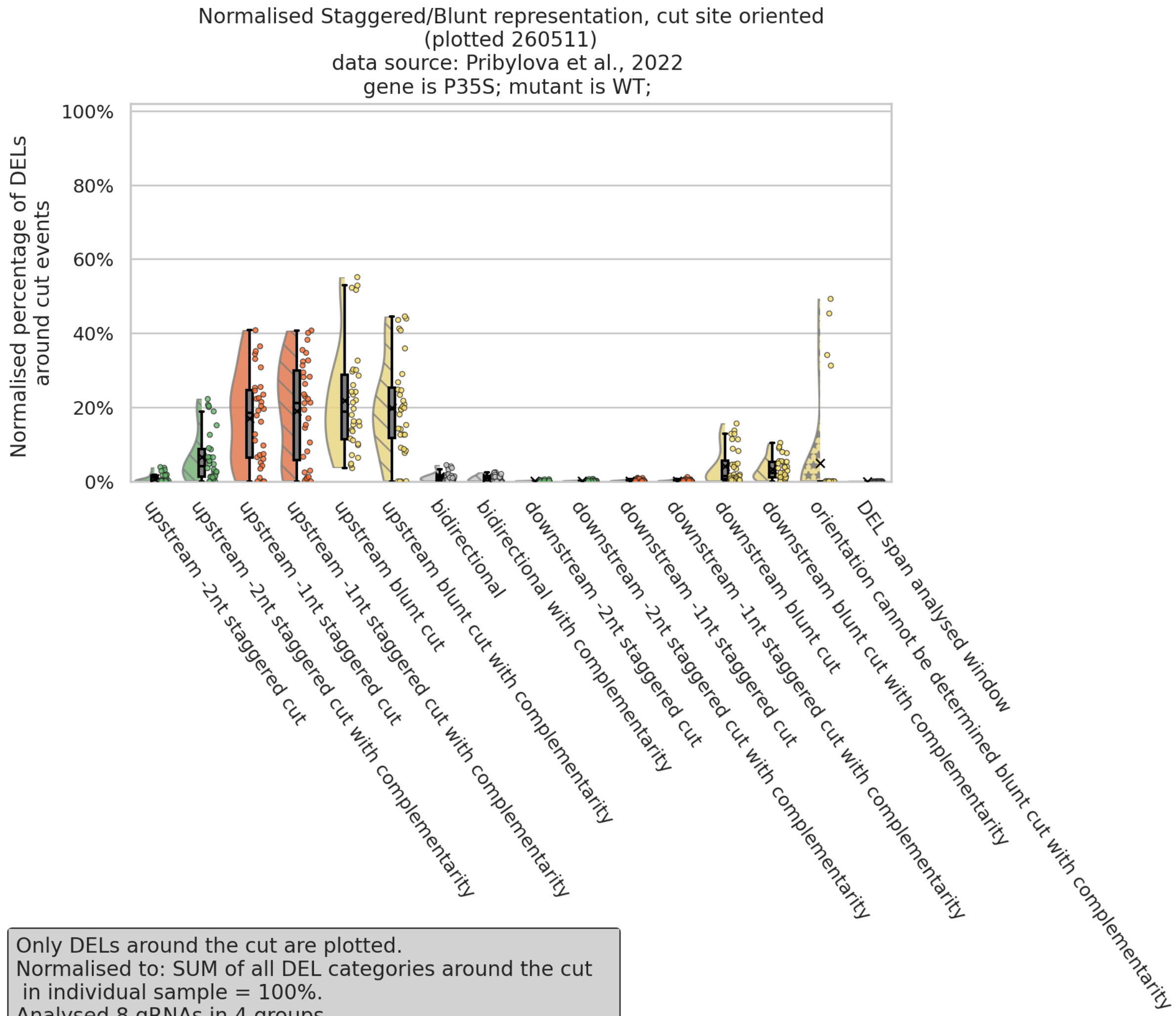


Only DELs around the cut are plotted.
Normalised to: SUM of all DEL categories around the cut
in individual sample = 100%.
Analysed 8 gRNAs in 4 groups.

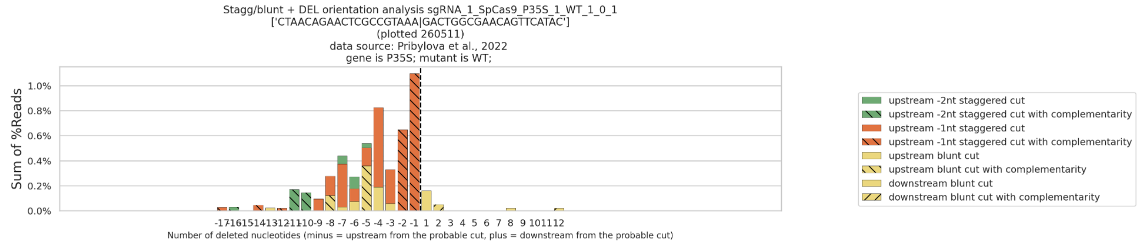


Data from deletions upstream and downstream from the probable cut only. Without bidirectional deletions.
Cannot be determined DELs were split between upstream and downstream.
Not normalised.

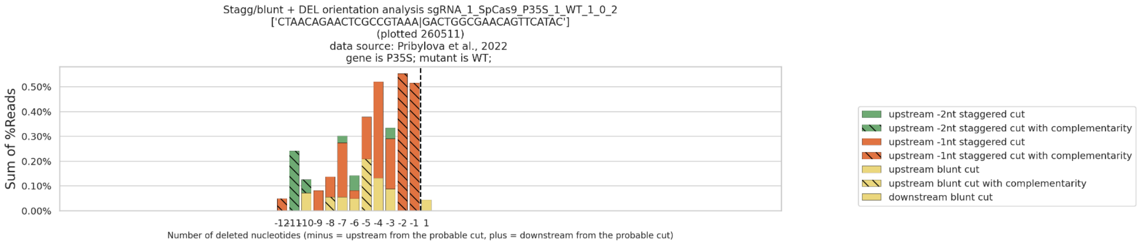


Data from deletions upstream and downstream from the probable cut only. Without bidirectional deletions.
Cannot be determined DELs were split between upstream and downstream.
Not normalised.

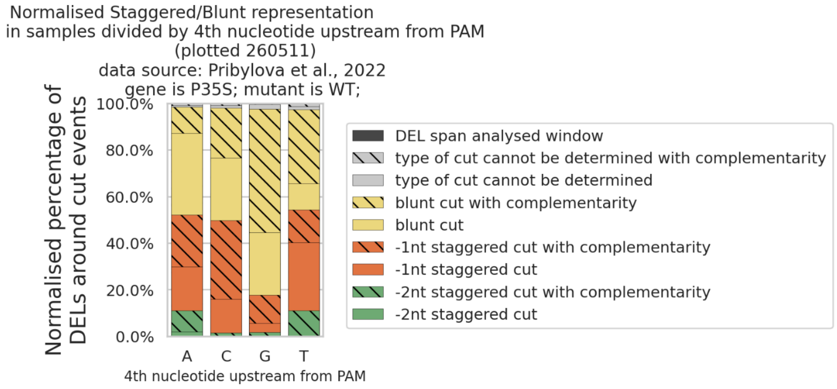


Only DELs around the cut are plotted.
Normalised to SUM of all DEL categories around the cut in individual sample = 100%,
then samples divided in 4th nt groups and mean was calculated for each DEL categry and plotted.
Analysed 8 gRNAs in 4 groups.

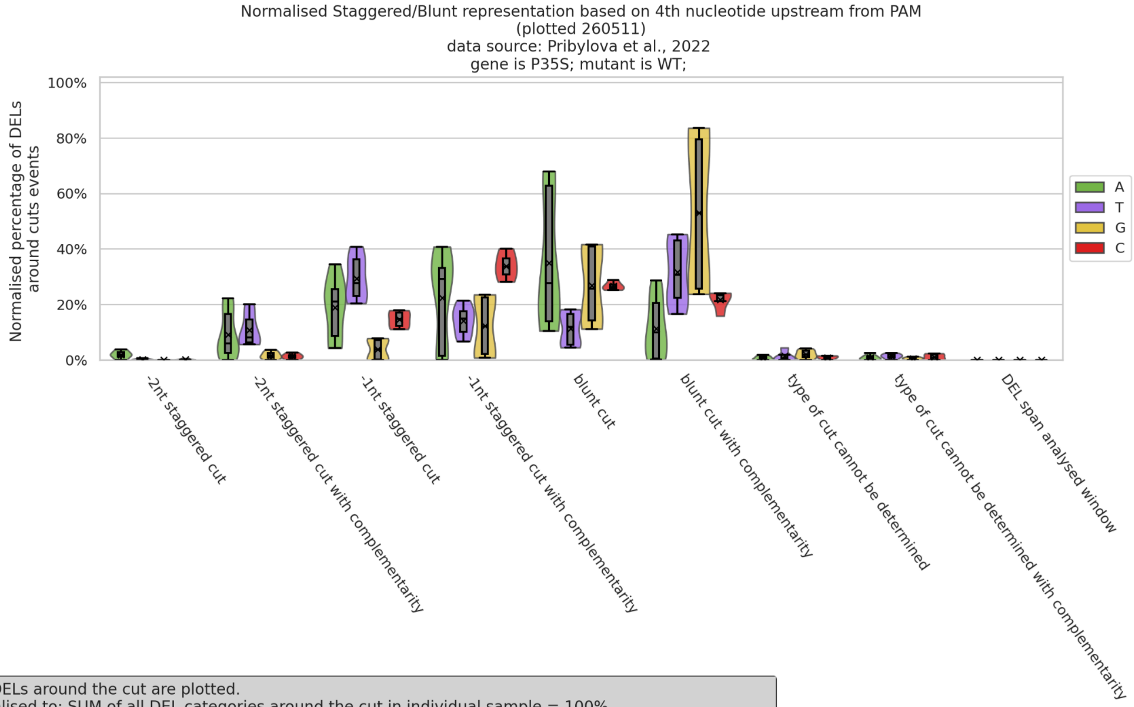


Only DELs around the cut are plotted.
Normalised to: SUM of all DEL categories around the cut in individual sample = 100%.
Analysed 8 gRNAs in 4 groups.

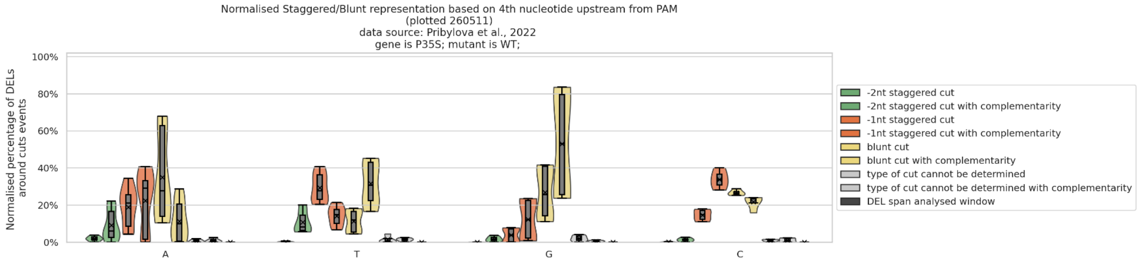


Only DELs around the cut are plotted.
Normalised to: SUM of all DEL categories around the cut in individual sample = 100%.
Analysed 8 gRNAs in 4 groups.

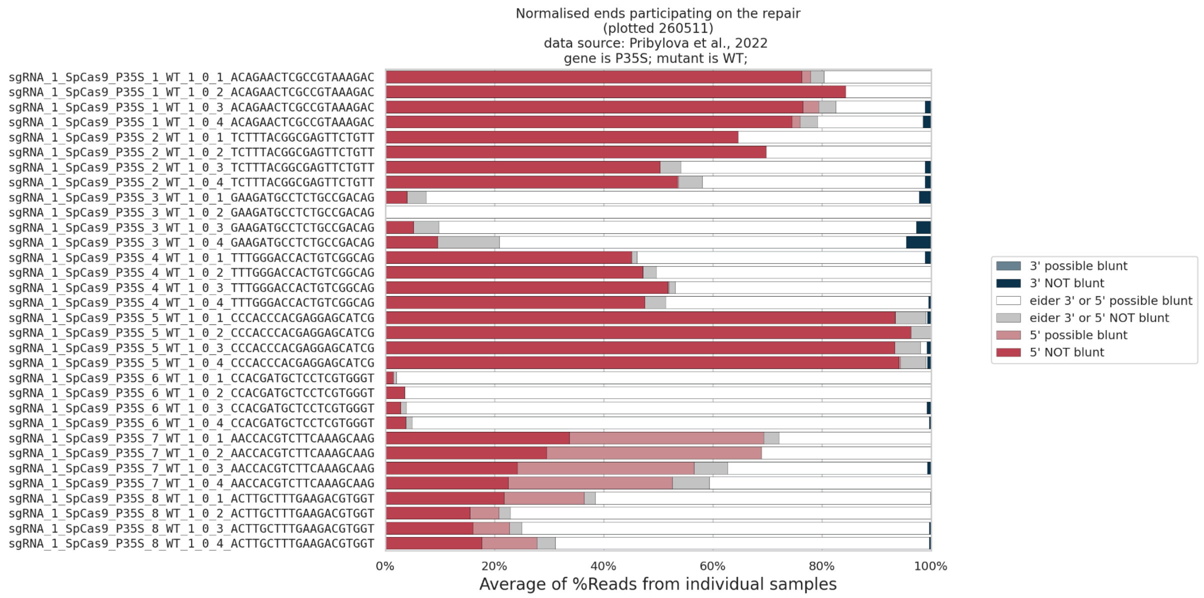


Only DELs with complementarity are plotted.
Normalised to: SUM of all ends in individual sample = 100%.
Analysed 8 gRNAs in 4 groups.

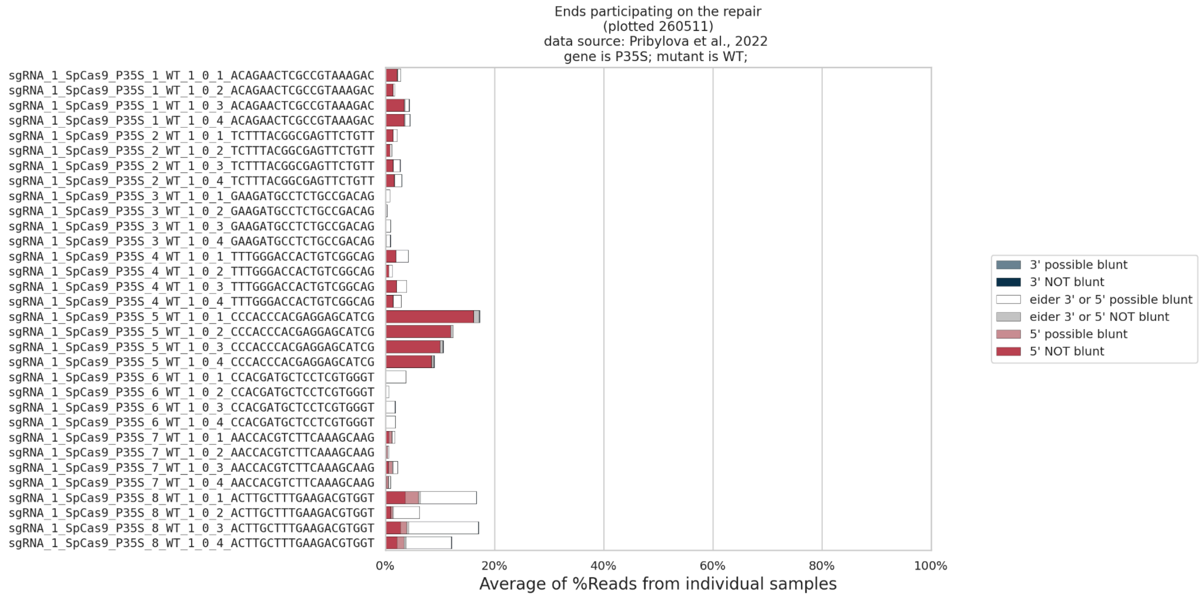


Only DELs with complementarity are plotted.
Analysed 8 gRNAs in 4 groups.

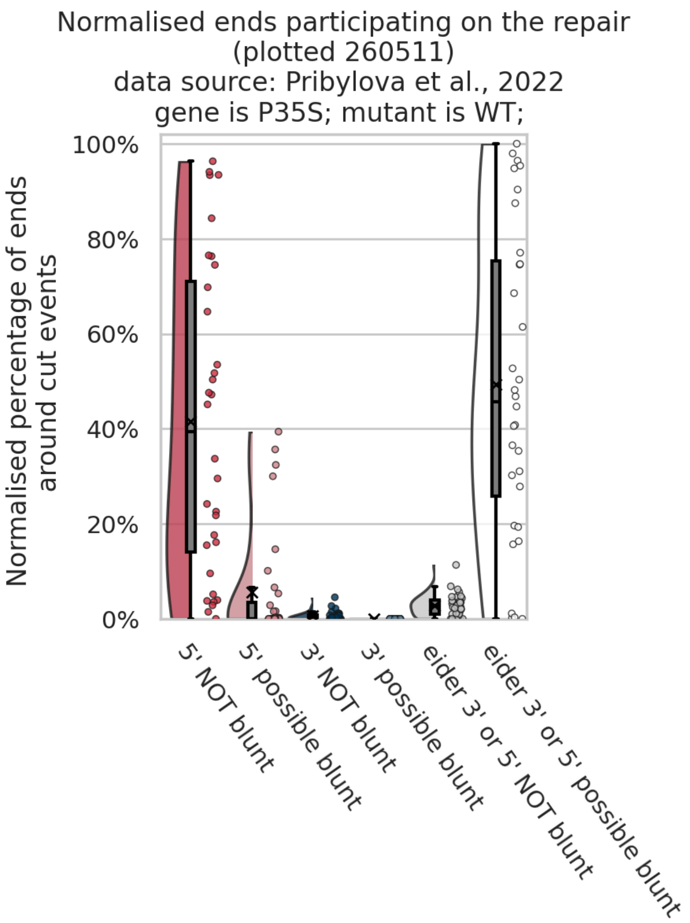


Only DELs with complementarity are plotted.
Normalised to: SUM of all ends in individual sample = 100%
Analysed 8 gRNAs in 4 groups.

Ends participating on the repair
(plotted 260511)
data source: Pribylova et al., 2022
gene is P35S; mutant is WT;

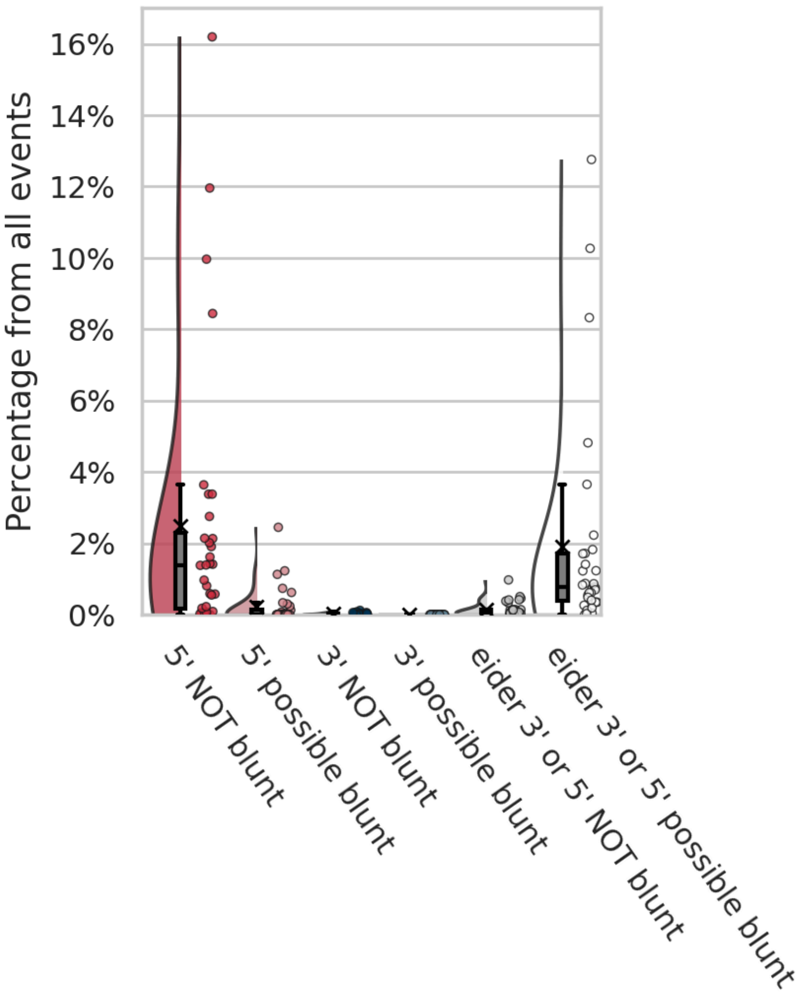


Only DELs with complementarity are plotted.
Analysed 8 gRNAs in 4 groups.

End participating on the repair + DEL orientation analysis sgRNA_1_SpCas9_P35S_1_WT_1_0_1
['CTAACAGAACTCGCCGTAAA|GACTGGCGAACAGTTCATAC']
(plotted 260511)
data source: Pribylova et al., 2022
gene is P35S; mutant is WT;

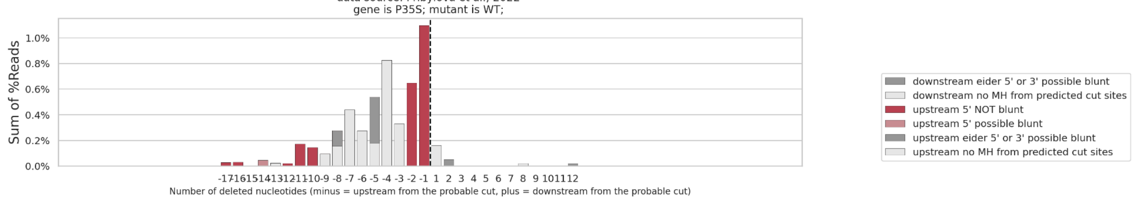


Data from deletions around cut only.
Without bidirectional deletions.
Cannot be determined was split between upstream and downstream.
Not normalised data.

End participating on the repair + DEL orientation analysis sgRNA_1_SpCas9_P35S_1_WT_1_0_2
['CTAACAGAACTCGCCGTAAA|GACTGGCGAACAGTTCATAC']
(plotted 260511)
data source: Pribylova et al., 2022
gene is P35S; mutant is WT;

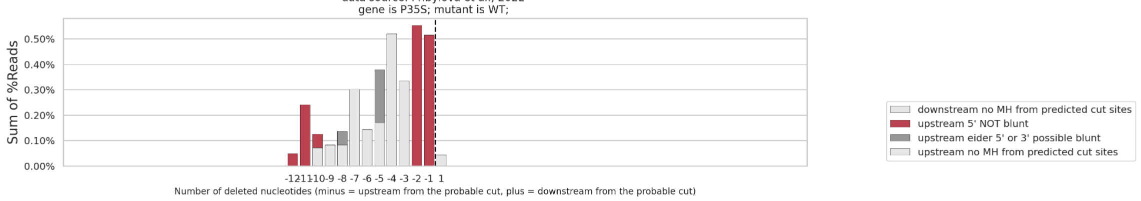


Data from deletions around cut only.
Without bidirectional deletions.
Cannot be determined was split between upstream and downstream.
Not normalised data.
